# TGF-β inhibitor accelerates BMP4-induced cochlear gap junction formation during *in vitro* differentiation of embryonic stem cells

**DOI:** 10.1101/2020.02.17.952481

**Authors:** Ichiro Fukunaga, Cheng Chen, Yoko Oe, Keiko Danzaki, Sayaka Ohta, Akito Koike, Ayumi Fujimoto, Katsuhisa Ikeda, Kazusaku Kamiya

## Abstract

Mutations in the connexin 26 (CX26)/gap junction beta-2 (*GJB2*) gene are the most frequent cause of hereditary deafness worldwide. Using mouse induced pluripotent stem cells (iPSCs) and a BMP4 signal-based floating and adherent culture system, we recently produced *in vitro* responsible for *GJB2*-related deafness (CX26-gap junction plaque-forming cells, CX26GJCs). However, to use these cells as a disease model platform for high-throughput drug screening or regenerative therapy, cell yields must be substantially increased. In addition to BMP4, presently uncharacterized factors may also induce CX26 gap junction (GJ) formation. A floating culture with embryonic stem cell (ESC) treatment and BMP4/TGF-β inhibitor (SB431542:SB) has been shown to result in greater production of isolatable CX26-positive small vesicles (CX26+ vesicles) and higher *Gjb2* mRNA levels than BMP4 treatment alone, suggesting that SB may promote BMP4-mediated production of CX26+ vesicles in a dose-dependent manner, thereby increasing the yield of highly purified CX26GJCs.

In the present study, we first demonstrated that SB accelerates BMP4 induced GJ formation during stem cell differentiation. By controlling the concentration and timing of SB supplementation with CX26+ vesicle purification, large-scale production of highly purified CX26GJCs suitable for high-throughput drug screening or regenerative therapy for *GJB2*-related deafness may be possible.

## Introduction

Hearing loss is the most common congenital sensory impairment worldwide[1]. Approximately 1 child in 1,000 is born with severe hearing loss or will develop hearing loss during early childhood, which is known as prelingual deafness[2, 3]. Approximately half of such cases are attributable to genetic causes[4]. Mutations in the gap junction beta-2 (*GJB2*) gene, which encodes connexin 26 (CX26), are the most common genetic cause of non-syndromic sensorineural hearing loss, accounting for ∼50% of such hearing loss in children[5, 6].

CX26 and CX30, encoded by *GJB6*, are expressed in non-sensory cochlear supporting cells and in such cochlear structures as the spiral limbus, stria vascularis, and spiral ligament[7–12]. In contrast, CXs are not expressed in hair cells[9, 11–13].

CX26 and CX30 have been shown to form functional heteromeric and heterotypic gap junction (GJ) channels in the cochlea[14] as well as in *in vitro* experiments[15]. At the plasma membrane, GJs further assemble into semi-crystalline arrays known as gap junction plaques (GJPs) containing tens to thousands of GJs[16].

GJs facilitate the rapid removal of K^+^ from the base of cochlear hair cells, resulting in cycling of K^+^ back into the endolymph of the cochlea to maintain cochlear homeostasis[17]. Several studies have reported that the deletion of *GJB2* can cause cochlear developmental disorders besides deafness, such as tunnel of Corti, Nuel’s space, or spaces surrounding the outer hair cells[18–23].

We previously showed that disruption of CX26-GJPs is associated with *Gjb2*-related hearing-loss pathogenesis and that assembly of cochlear GJPs is dependent on CX26[24]. Thereafter, we showed that the transfer of *Gjb2* into the cochleae using an adeno-associated virus significantly improved GJP formation and auditory functions in a mouse model[25].

Furthermore, we have recently reported that induced pluripotent stem cells (iPSCs)-derived functional CX26-GJP-forming cells (CX26GJCs), as found in the cochlea supporting cells[26], which is unlike previous studies that targeted the generation of cochlear hair cells from embryonic stem cells (ESCs) and iPSCs[27–39].

Thus, CX26GJC could be generated from a floating culture (SFEBq culture; a serum-free floating culture of embryoid body-like aggregates with quick reaggregation) followed by an adherent culture[26].

SFEBq culture is superior for generating ectoderm-derived tissues, which can give rise to forebrain, midbrain, adenohypophysis, retinal tissue, and otic sensory epithelia[29, 38, 40–45]. Furthermore, in SFEBq culture, ectodermal tissues develop in an epithelial form similar to that in *in vivo* counterparts.

However, to use these cells as a disease model for drug screening or other large-scale assays, the cell culture system must be improved to increase the number of cells available at a single time. Our previous research suggested that CX26 expressing vesicles formed in day 7 aggregate are the origin of CX26-GJP-forming cells in the 2D culture[26]. If CX26 small vesicles in SFEBq culture from ES/iPS cells could be obtained in a substantial quantity, we could obtain an adequate number of CX26-GJP-forming cells in the 2D culture.

Several studies have shown that BMP4 signaling plays a key role in embryonic development[46–50] and *in vitro* differentiation of ESCs/iPSCs[51–53].

In SFEBq culture, BMP4 upregulated a non-neural ectoderm marker (*Dlx3*) and downregulated a neuroectoderm marker (*Sox1*)[29]. Furthermore, BMP4 drives CX expression and CX-mediated cell-to-cell communication[54–56].

Similarly, TGF-β inhibitor (SB431542:SB) has been implicated in efficient neural conversion of ESCs and iPSCs via inhibition of SMAD signaling[57, 58], and by blocking the progression of stem cell differentiation toward trophectoderm, mesoderm, and endoderm lineages[59]. In SFEBq culture using mouse ESCs, SB inhibition of TGF-β signaling is thought to promote proper non-neural induction following BMP4 treatment[29, 60]. Although cochlear CX26-GJP-forming cells in cochlear non-sensory regions should be part of the non-neural ectoderm, it has not been determined whether SB can accelerate BMP4-induced CX expression or GJ formation.

Given this background, we hypothesized that SB plays a key role in the differentiation of CX26GJCs. Therefore, in the present study, we evaluated modified SFEBq culture conditions incorporating BMP4 and/or SB with the aim of generating CX26GJCs from mouse ESCs with a greater potential to differentiate than that of iPSCs. The large-scale production of CX26-GJP-forming cells could be used for high-throughput drug screening related to deafness induced by mutations in *GJB2*.

## Materials and methods

All the experimental protocols using mouse tissues were approved by the Institutional Animal Care and Use Committee at Juntendo University School of Medicine and were conducted in accordance with the US National Institutes of Health Guidelines for the Care and Use of Laboratory Animals. Adult mice (10-weeks-old) were obtained from CLEA Japan, Inc. All methods were carried out in accordance with relevant guidelines and regulations.

### ESC culture

Mouse ESCs (EB5 cells)[61, 62] were provided by the RIKEN Bio Resource Center Cell Bank and maintained under feeder-free conditions with 2i-LIF medium[63]. Briefly, ESCs were maintained on gelatin containing N2B27 medium consisting of a 1:1 (v/v) mixture of Advanced DMEM/F12 and neurobasal medium (Invitrogen) supplemented with 1 mM GlutaMAX (Invitrogen), 1% N2 supplement (Invitrogen), 2% B27 supplement (Invitrogen), 3 μM CHIR99021 (Stemgent), 1 μM PD0325901 (Santa Cruz), and 1,000 U ml^−1^ of leukemia inhibitory factor (Millipore).

### Differentiation of ESCs

Induction of CX26GJCs was performed as shown in Fig 1A. Briefly, ESCs were dissociated with Accutase (Innovative Cell Technologies, Inc.), suspended in differentiation medium (G-MEM, Gibco) supplemented with 1.5% (v/v) knockout serum replacement (Gibco), 0.1 mM nonessential amino acids (Gibco), 1 mM sodium pyruvate (Gibco), and 0.1 mM 2-mercaptoethanol), and then plated at 100 μl per well (3,000 cells) in 96-well low-cell attachment V-bottom plates (Sumitomo Bakelite). Recombinant BMP4 was obtained from Miltenyi Biotec and SB431542 was obtained from Tocris Bioscience. On day 1, half of the medium (50 μl) per well was replaced with fresh differentiation medium containing 4% (v/v) Matrigel (BD Bioscience). On day 3, one of three types of media was added to the culture: medium containing BMP4 (10 ng/ml, final concentration), SB (1–10 μM, final concentration), or both factors at the aforementioned concentrations.

**Fig 1.**
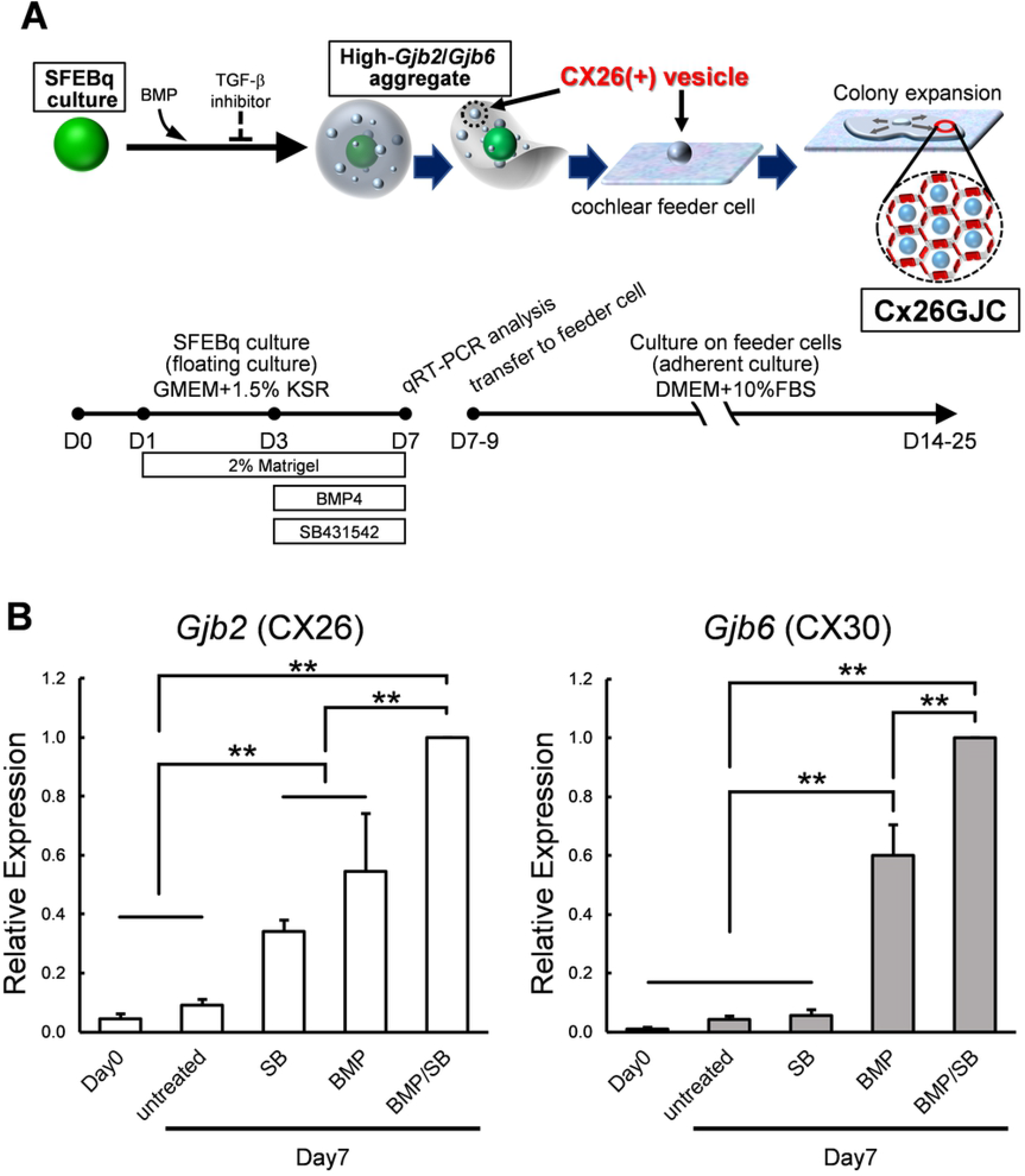
Culture conditions for cells expressing high levels of *Gjb2* (CX26) and *Gjb6* (CX30) mRNA. **(A):** A schematic procedure for differentiating Connexin26 gap junctional plaque forming cells (Cx26GJCs) from mouse ESCs. SFEBq; serum-free floating culture of embryoid body-like aggregates with quick reaggregation, KSR; knockout serum replacement, SB431542; TGF-beta inhibitor. **(B):** Relative expression of mRNA at day 0 (for undifferentiated ES cells) and at day 7 for untreated, BMP4-treated, SB-treated, and BMP4/SB-treated aggregates. The mRNA expression levels were normalized to those of BMP4 culture on day 7. (qRT-PCR: n = 5. For assessments, the procedures were repeated five times to generate cells. Each experiment used 8 aggregates. Differences between samples were assessed by One-way ANOVA and multiple comparison test; **, p < 0.01. The data are expressed as mean ±standard error.

BMP4 and SB stock solutions were prepared at 5× concentration in fresh medium. On days 7–11, the aggregates were semi-dissected and the small vesicles were mechanically isolated and collected using forceps. The small vesicles were transferred into adherent culture containing TRICs in the growth medium, which was composed of DMEM GlutaMAX (Gibco) and 10% (w/v) FBS.

TRICs were generated by exposing cochlear tissue to trypsin and screening for trypsin-resistant cells. The mouse cochlear tissue (10-weeks-old) used for preparation of the TRICs included the organ of Corti, basilar membrane, and lateral wall, and mainly comprised supporting cells, hair cells, cochlear fibrocytes, and other cells in the basilar membrane. This cell line was used as inner-ear derived feeder cells to proliferate the otic progenitor cells. For the feeder cell layer, 3 × 10^5^/cm^2^ TRICs were seeded into gelatin-coated wells of 24-well culture plates after mitomycin C (10 mg/ml) treatment for 3 h.

### qRT-PCR of *Gjb2* and *Gjb6* mRNA expression

Total RNA was isolated using reagents from an RNeasy Plus Mini kit (Qiagen) and reverse transcribed into cDNA using reagents from a Prime Script II first strand cDNA synthesis kit (Takara). Real-time PCR was performed with the reverse transcription products, TaqMan Fast Advanced Master Mix reagents (Applied Biosystems), and a gene-specific TaqMan Probe (see below; Applied Biosystems) on a StepOne Real-Time PCR system (Applied Biosystems). Each sample was run in triplicate. Applied Biosystems StepOne software was used to analyze the Ct values of the different mRNAs normalized to expression of the endogenous control, actin beta mRNA. TaqMan Probes (Assay ID; Applied Biosystems) were used to detect the expression of mouse *Gjb2* (Mm00433643_s1), *Gjb6* (Mm00433661_s1), and *actin beta* mRNAs (Mm02619580_g1).

### Immunostaining and image acquisition

Aggregates were fixed with 4% (w/v) paraformaldehyde in 0.01 M phosphate-buffered saline (PBS) for 1 h at room temperature. For whole mounts, the aggregates were permeabilized with 0.5% (w/v) Triton X-100 (Sigma-Aldrich) in 0.01 M PBS for 30 min. Then, the samples were washed twice with 0.01 M PBS and blocked with 2% (w/v) bovine serum albumin in 0.01 M PBS for 30 min.

Cells from adherent cultures were fixed with 4% (w/v) paraformaldehyde in 0.01 M PBS for 15 min at room temperature and, then, permeabilized with 0.5% (w/v) Triton X-100 in 0.01 M PBS for 5 min. Samples were washed twice with 0.01 M PBS and blocked with 2% (w/v) bovine serum albumin in 0.01 M PBS for 30 min. For immunofluorescence staining, 1% (w/v) bovine serum albumin in 0.01 M PBS was used to dilute the primary and secondary antibody solutions. Each sample was incubated in a primary antibody solution—CX26 or CX30 (mouse IgG, 33-5800 or rabbit IgG, 71-2200, respectively, Life Technologies)—for 1 h after blocking. The secondary antibodies were Alexa Fluor 594–conjugated anti-mouse IgG (Invitrogen, A11032), Alexa Fluor 488–conjugated anti-rabbit IgG (Invitrogen, A11070), and phalloidin FITC (Invitrogen, A12379). Samples were washed twice with 0.01 M PBS and mounted with mounting medium (VECTASHIELD Mounting Medium with DAPI, Vector).

Fluorescence confocal images were obtained with an LSM510-META confocal microscope (Zeiss). Z-stacks of images were collected at 0.5 μm intervals, and the single-image stacks were constructed using LSM Image Browser (Zeiss). Three-dimensional images were constructed with z-stacked confocal images using IMARIS (Bitplane).

### Statistical analyses

The data were analyzed using Microsoft Excel software and are presented as the mean ± standard error. A two-tailed Student’s *t*-test, with a significance criterion of p < 0.05, was used to compare the GJP lengths. One-way ANOVA and multiple comparison test, with a significance criterion of p < 0.05, was used to compare *Gjb2* and *Gjb6* mRNA levels and the number of CX26+ vesicles.

## Results

### SB431542, an inhibitor of TGF-β signaling, promoted BMP4-induced *Gjb2/Gjb6* mRNA expression in SFEBq culture

CX26GJCs were induced from mouse ESCs using previously reported method[26], and the conditions required for differentiation were then assessed. ESCs were cultured in SFEBq medium containing BMP4, SB, or BMP4 plus SB. Aggregates were collected on day 7 and mRNA (*Gjb2* and *Gjb6*) levels in the different culture groups were measured. BMP4 and BMP4/SB treatments produced more *Gjb2/Gjb6* mRNA than SB and control cultures (Fig 1B). BMP4 is a factor that induces *Gjb2*/*Gjb6* mRNA expression during iPSC differentiation[26]. In addition, ESCs cultured in differentiation medium supplemented with BMP4 and SB showed greater expression levels of mRNA (*Gjb2*, 1.8-fold greater; *Gjb6*, 1.7-fold greater) compared with the BMP4 alone group.

### SB431542 promoted formation of CX26-expressing small vesicles in SFEBq culture

By day 7 of differentiation, the aggregates showed differentiated outer regions with a morphology similar to that reported previously[26]. Clear outer epithelia and small vesicles were observed beneath the outer epithelium of BMP4/SB-treated cells. By contrast, no small vesicles were observed for the control or SB-treated cells (Fig 2A, left column). To determine the location of CX26 in the cell aggregates, immunohistochemistry was performed. In BMP4 or BMP4/SB-treated aggregates, small vesicles containing CX26-GJP (hereafter called CX26+ vesicles) were observed (Fig 2A, right column). The aggregates were collected, and the numbers of CX26+ vesicles were compared among the different treatment groups (Fig 2B).

**Fig 2.**
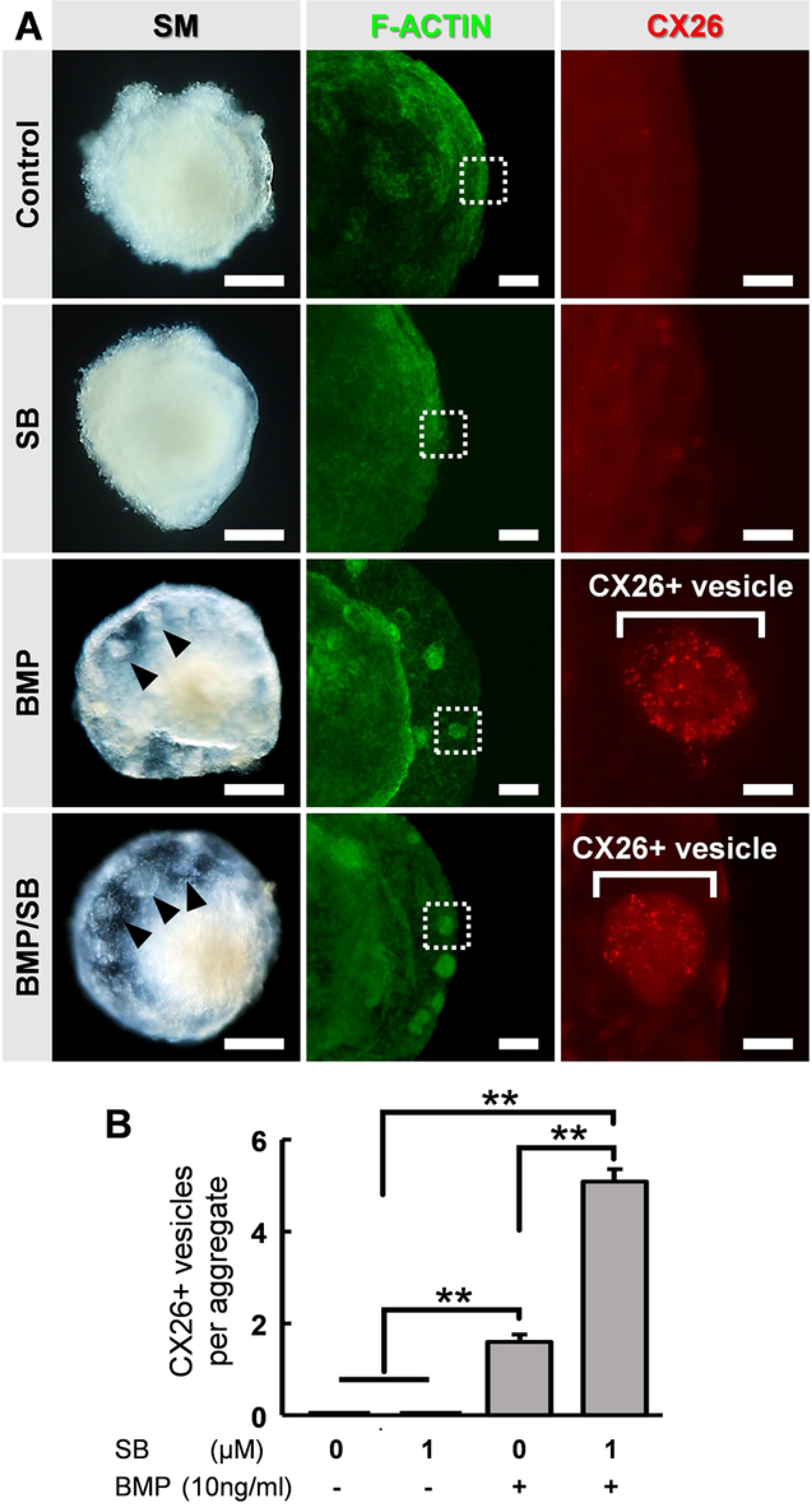
Stereomicroscopic images and immunostained aggregates derived from ESCs present on day 7. **(A):** Left column: stereomicroscopic images of cells on day 7. Middle column: F-actin immunostained cells (green). Right column: magnification of boxed regions in the middle columns. Immunostained for CX26 (red). Arrowheads point to CX26+ vesicles. Scale bars, 200 μm (left column); 100 μm (middle column); 20 μm (right column). **(B):** The average number of CX26+ vesicles per aggregate from SB-, BMP4-, and BMP4/SB-treated, and untreated control cultures (n = 24 aggregates. For assessments, the procedures were repeated three times to generate cells. Statistical differences between samples were assessed by One-way ANOVA and multiple comparison test; **, p < 0.01. The data are expressed as mean ± standard error.

Cells treated with BMP4/SB had more CX26+ vesicles (mean = 5.08 ± 0.28) than cells cultured only with BMP4 (mean = 1.6 ± 0.16). CX26+ vesicles were found to exist separately from the BMP4/SB-treated aggregates (Fig 3A and 3B), suggesting that they could be easily isolated. Numerous CX26+ vesicles were mechanically collected as a purified CX26GJC population (Fig 3C).

**Fig 3.**
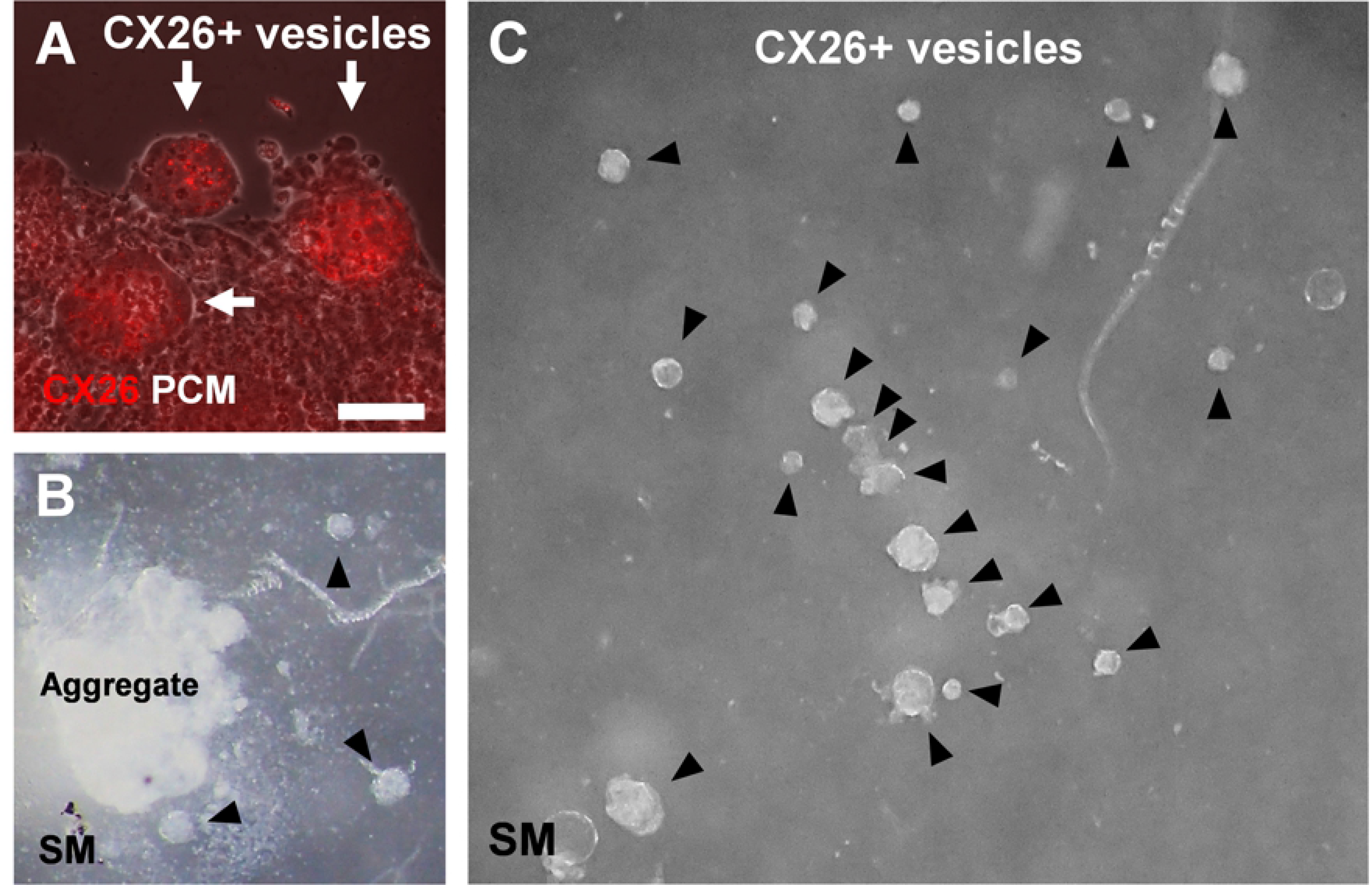
CX26+ vesicles in aggregates with BMP4 and SB supplementation. **(A):** Merged images of CX26-immunostained (red) and phase-contrast microscopy-imaged (PCM, white) ESC aggregates at day 7. Arrows point to CX26+ vesicles containing CX26GJCs. The scale bar represents 50 μm. **(B-C):** Stereomicroscopic (SM) images of isolated CX26+ vesicles from ESC aggregates at day 7. Arrowheads point to CX26+ vesicles. CX26+ vesicles were easily isolated from semi-dissected ESC aggregates **(B)** and then mechanically collected **(C)**.

In the confocal analysis of the BMP4/SB-treated day-7 aggregates, CX26-expressing cells were disseminated throughout the numerous CX26+ vesicles (Fig 4A and S1 Video). These cells formed CX26-positive GJs at their cell-cell borders (Fig 4B). In the 3D construction of the confocal images, we observed large planar CX26-containing GJPs (Fig 4C and S2 Video) which, as we reported previously[24, 26], are characteristic of mouse cochlea.

**Fig 4.**
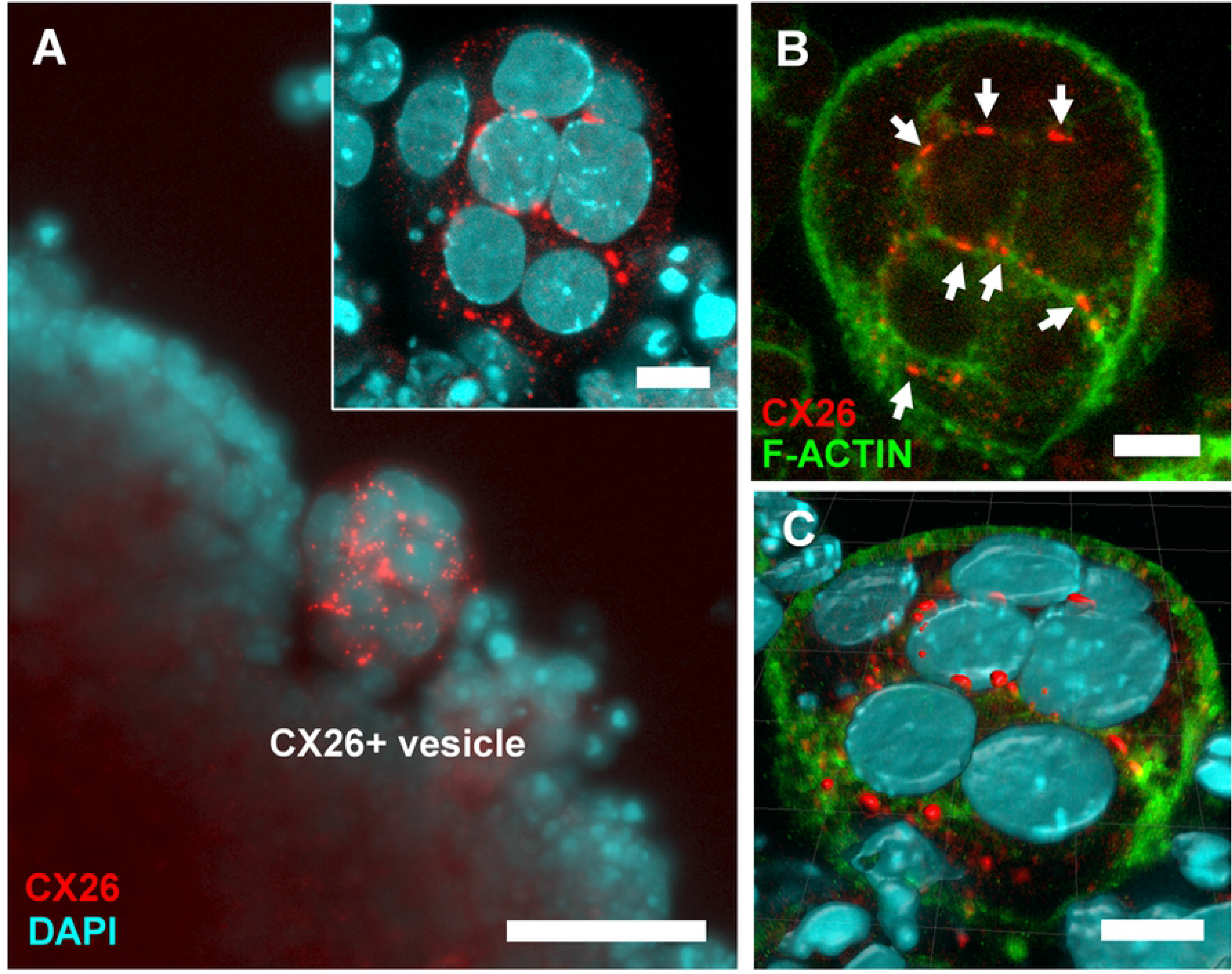
Confocal images of CX26+ vesicles in BMP4/SB-treated ESC aggregates. **(A):** Merged image of CX26-immunostained (red) and DAPI-stained (blue) cells in CX26+ vesicles. (inset in A) Magnification of a CX26+ vesicle. **(B):** Merged images of CX26-immunostainded (red) and F-actin stained (green) cells in the same region as in **(A)**. Arrows point to GJPs. **(C):** The 3D image was reconstructed from the same region as shown in (inset in A, B). Scale bars: 50 μm (A), 10 μm: (inset in A, B, and C).

### ESC-derived CX26GJC co-expressed CX30 in adherent cultures formed GJPs

Between day 7 and 9, BMP4/SB-treated aggregates were transferred onto cochlear-derived feeder cells, namely trypsin-resistant inner-ear cells (TRICs), as follows. The differentiated regions with CX26+ vesicles were separated from the day 7 aggregates and subcultured in Dulbecco’s modified Eagle’s medium (DMEM) GlutaMAX, 10% (v/v) fetal bovine serum (FBS) on TRIC feeder cells. The subcultured regions containing CX26+ vesicles colonized the TRIC feeder cells. In the adherent culture at day 15, CX26-containing GJPs were preserved (Fig 5A–C), as found in cochlear supporting cells. The mean length of the longest dimension of the GJPs along a single cell border was 1.91 ± 0.11 μm for BMP4/SB-treated aggregates, which was significantly increased to 5.39 ± 0.25 μm in the adherent culture on TRIC feeder cells (Fig 5D). To assess the similarities between these cells and cochlear cells, we characterized the expression of CX30, which is frequently absent in hereditary deafness. CX30 co-localized with CX26 in most CX26-GJPs in the differentiated cells (Fig 5E– H), suggesting that CX26 and CX30 were the two main components of these GJPs, as was found for cochlear cells[26].

**Fig 5.**
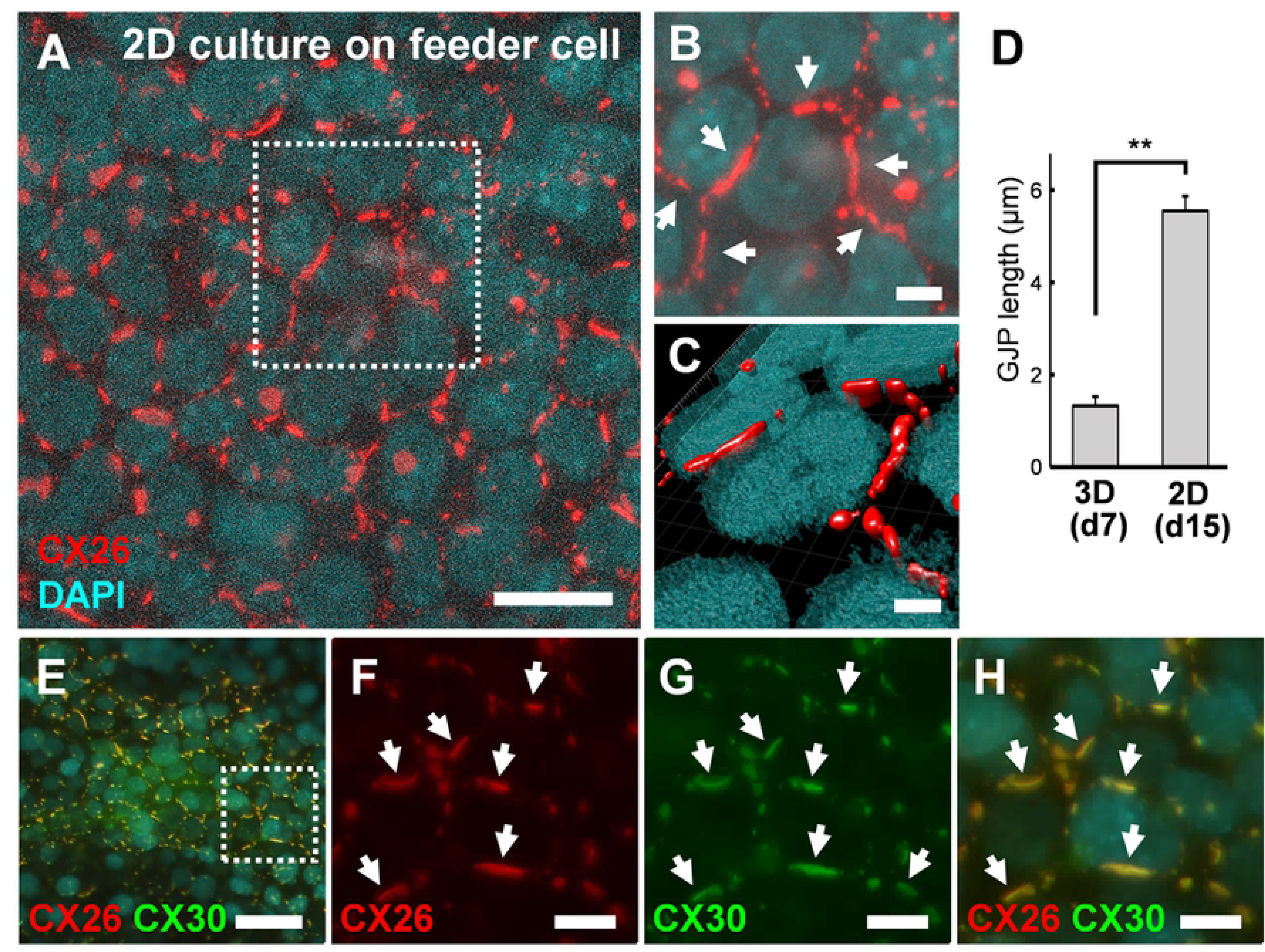
ESC-derived CX26GJC formed large CX26GJPs and CX26/CX30-containing GJPs. **(A):** Merged images of CX26-immunostained (red) and DAPI-stained (blue) cells from an adherent culture at day 15. **(B):** Magnification of the boxed region in **(A)**. Arrows point to GJPs. **(C):** The 3D image was reconstructed from the same region as shown in **(B)**. **(D):** Mean lengths of the longest dimension of the GJPs along a single cell border in SFEBq culture (3D) at day 7 and adherent culture (2D) at day 15. (SFEBq culture, n = 43 cell borders from 5 aggregates; adherent culture, n = 41 cell borders from 4 wells. For assessments, the procedures were repeated three times to generate cells. Statistical differences between samples were assessed by Student’s *t*-test, **, p < 0.01. The data are expressed as mean ± standard error. **(E):** Merged images of CX26-immunostained (red), CX30-immunostained (green), and DAPI-stained (blue) cells from the adherent culture at day 15. **(F-H):** Magnification of the boxed region in **(E)**. Staining for CX26 (red), CX30 (green), and DAPI (blue) was as in **(E)**. Arrows point to the large GJPs. Scale bars: 20 μm (A and E), 10 μm (F-H), 5 μm (B), and 3 μm (C).

### The amount of mRNA expression and the number of CX26 positive small vesicles were increased by SB431542 addition in a dose-dependent manner

Finally, to produce a large number of CX26GJC in SFEBq culture, we examined whether the differentiation from ES cells to CX26GJC depends on the concentration of SB431542.

As a result of qRT-PCR analysis, in BMP4 and SB combination, 5 μM and 10 μM, SB-treated aggregates showed greater expression level of *Gjb2* mRNA (BMP4/SB 5 μM, 1.7-fold greater; BMP4/SB 10 μM, 1.7-fold greater) compared with the BMP4/SB 1 μM. The expression level of *Gjb6* did not differ depending on the concentration of SB (Fig 6A). In the SB alone group, there was no difference in the expression level of *Gjb2*/*Gjb6* depending on the concentration of SB (S1A Fig). Thereafter, we determined the number of CX26 positive vesicles in day 7 aggregates by immunohistochemistry. In BMP4 and SB combination, 5 μM and 10 μM, SB-treated aggregates showed greater numbers of CX26+ vesicles (BMP4/SB 5μM, 1.5-fold greater; BMP4/SB 10μM, 1.3-fold greater) compared with the BMP4/SB 1μM (Fig 6B). Conversely, in the SB alone group, there was no difference in the number of small vesicles depending on the concentration of SB (S1B Fig).

**Fig 6.**
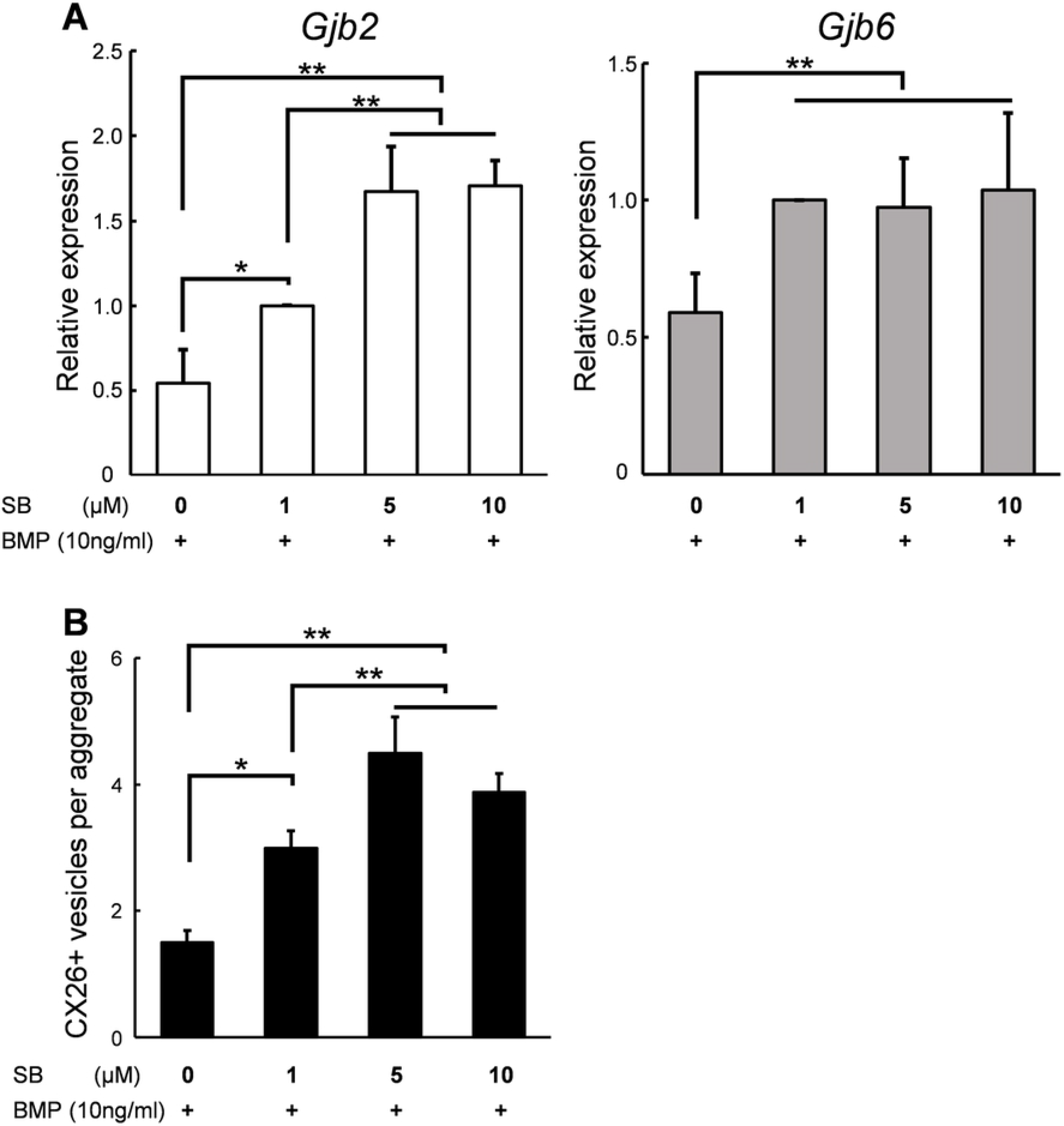
Dose-dependent manner of SB concentration in SFEBq culture expression level of mRNA and the number of CX26 + vesicles. **(A):** Relative expression of mRNA at day 7 for BMP4-treated (SB 0 μM + BMP4 10 ng/ml), and BMP4/SB (SB 1–10 μM + BMP4 10 ng/ml) treated aggregates. The mRNA expression levels were normalized to those of BMP4/SB 1 μM (SB 1 μM + BMP4 10 ng/ml) culture on day 7. (qRT-PCR, n = 5). For assessments, the procedures were repeated five times to generate cells. Each experiment used 8 aggregates. Differences between samples were assessed by the Scheffe multiple comparison test; **, p < 0.01. The data are expressed as mean ± standard error. **(B):** The average number of CX26+ vesicles per aggregate from BMP4-treated (SB 0 μM + BMP4 10 ng/ml), and BMP4/SB (SB 1–10 μM + BMP4 10 ng/ml) treated aggregates. (n = 9 aggregates). For assessments, the procedures were repeated three times to generate cells. Statistical differences between samples were assessed by One-way ANOVA and multiple comparison test: *, p < 0.05; **, p < 0.01. The data are expressed as mean ± standard error.

## Discussion

Several previous studies have reported BMP4 signaling-induced stem cell differentiation[29, 51–53], differentiation into CX43-expressing cells as cardiomyocytes[64], and CX43 expression in mouse embryonal cells[54–56, 65]. Although SB is reportedly involved in stem cell differentiation[29, 58–60, 66], it has not been shown to promote CX expression or GJ formation during stem cell differentiation or in mature cells. For the large-scale production of CX26GJCs, we evaluated the necessary conditions for the differentiation of pluripotent stem cells using modified SFEBq culture containing BMP4 and/or SB with ESC, which has a more stable differentiation potential than iPSCs.

We found that BMP4/SB treatment resulted in significantly greater production of CX26+ vesicles (Fig 2A and 2B) and significantly greater amount of *Gjb2* and *Gjb6* mRNAs (Fig 1B) than treatment with only BMP4 (Fig 7). Thus, these results suggest that the increase in CX26+ vesicles is because of the upregulation of mRNA (*Gjb2* and *Gjb6*) and SB strongly promoted BMP4-mediated differentiation of CX26GJCs (Fig 7).

**Fig 7.**
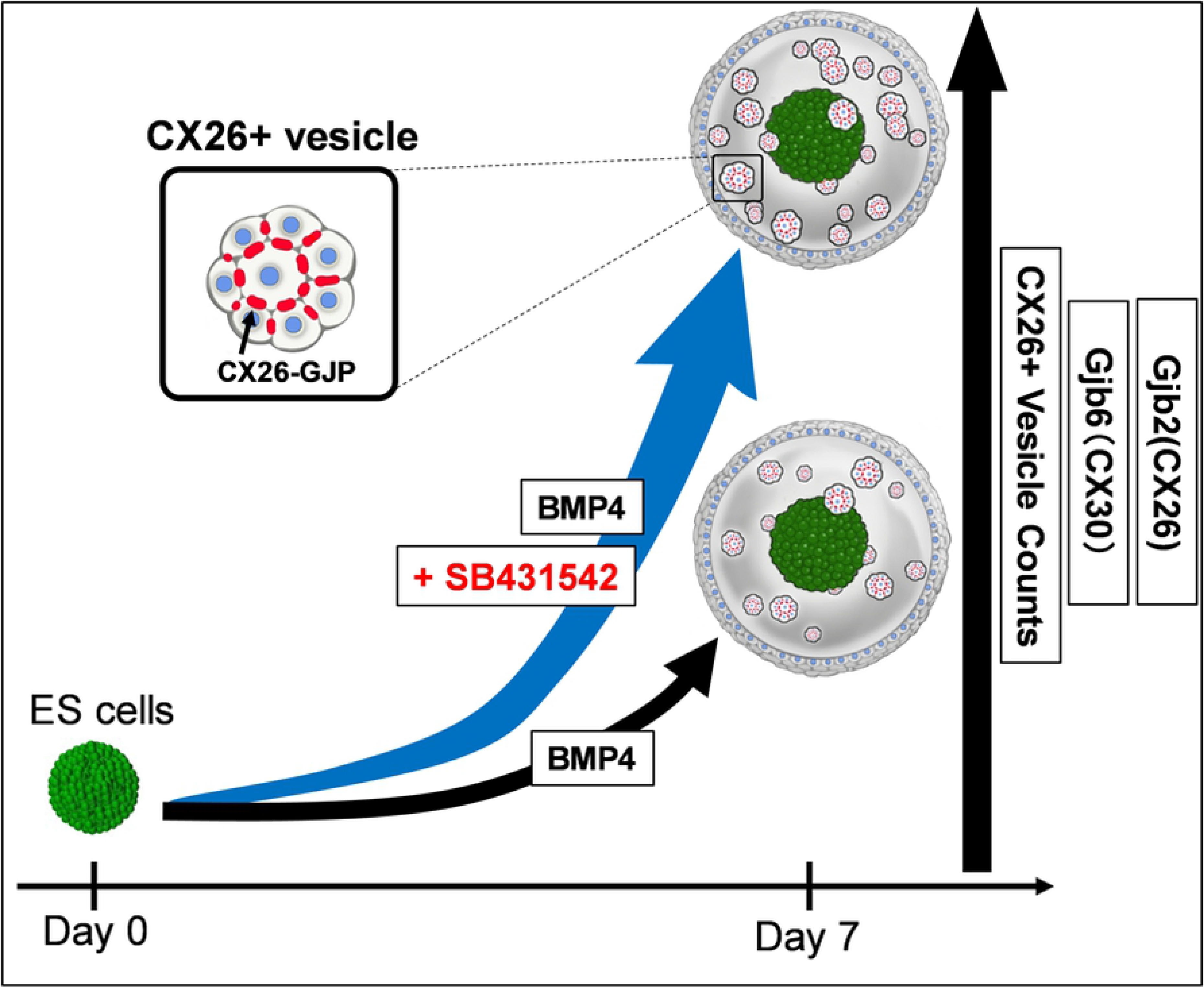
A schematic illustration of the effect of SB431542 on CX26 GJ formation in BMP4-induced ESC differentiation. In the BMP4-based inner-ear 3D differentiation from ESCs, SB431542 supplementation was associated with significantly higher mRNA levels of *Gjb2* (CX26) and *Gjb6* (CX30) and CX26+ vesicle counts than under SB431542 absent condition. SB431542 was demonstrated to be an accelerator of gap junction formation.

In the adherent culture that included TRIC feeder cells, we observed proliferation of CX26GJCs (Fig 5A) and long CX26-containing GJPs (Fig 5D), similar to observations when iPSCs were used[26]. Because the differentiated, aggregated cells co-expressed CX30 and CX26 (Fig 5E–H), it is likely that some cells were cochlear non-sensory cells containing CX26/CX30 GJPs that proliferated on the cochlear feeder cells after isolation of CX26+ vesicles.

When BMP4 was present in the SFEBq culture, the addition of SB appeared to promote BMP4-mediated formation of CX26+ vesicles. After transferring the CX26+ vesicles onto cochlear feeder cells, the lengths of the GJPs of the proliferated CX26-expressing cells increased and the GJPs were observed to contain CX26/CX30. These results indicate that the number of CX26GJCs increased in adherent culture due to the increase in CX26+ vesicles by SB treatment. Therefore, we suggest that SB promotes BMP4-mediated formation of CX26+ vesicles in SFEBq culture. Mechanical dissection of the aggregates (Fig 3B and 3C) suggested that CX26+ vesicles could be easily purified and isolated for large-scale production of CX26GJCs after adherent culture on feeder cells (Fig 5).

Furthermore, in the combination of BMP4/SB in SFEBq culture, increases in mRNA expression and CX26 positive small vesicles were found to depend on the concentration of SB. However, in the expression level of *Gjb2* and the number of CX26 positive vesicles, there was no significant difference between 5 μM and 10 μM SB addition (Fig 6). These results indicated that CX26GJCs could be most efficiently induced in the BMP4/SB 5 μM combination.

These data suggest that SB promotes BMP4-mediated production of CX26+ vesicles in a dose-dependent manner, thereby increasing the yield of highly purified CX26GJCs.

This is the first study to show that SB431542 accelerates BMP4-induced GJ formation during *in vitro* differentiation of ESCs (Fig 7).

By controlling the timing and concentration of SB with CX26+ vesicle purification, large-scale production of highly purified CX26GJCs for high-throughput screening of drugs that target *GJB2*-related deafness should be possible.

## Acknowledgments

Kazusaku Kamiya: Conceptualization, Data Curation, Formal Analysis, Funding Acquisition, Investigation, Methodology, Project Administration, Resources, Supervision, Validation, Visualization, Writing – Original Draft Preparation, Writing – Review & Editing

Ichiro Fukunaga: Data Curation, Formal Analysis, Funding Acquisition, Investigation, Methodology, Resources, Validation, Visualization, Writing – Original Draft Preparation, Writing – Review & Editing

Cheng Chen, Yoko Oe, Keiko Danzaki, Sayaka Ohta, Akito Koike, Ayumi Fujimoto: Resources, Validation, Katsuhisa Ikeda: Conceptualization, Resources

## Supporting information

**S1 Fig. Effects of the addition of various concentrations of SB on the expression level of mRNA and the number of CX26 + vesicles.** (A): Relative expression of mRNA at day 7 for SB-treated (SB 1-10μM), and BMP4/SB (SB 1μM+BMP4 10ng/ml) treated aggregates. The mRNA expression levels were normalized to that of BMP4/SB 1μM (SB 1μM+BMP4 10ng/ml) culture on day 7. (qRT-PCR: n = 5. For assessments, five time procedures were repeated to generate cells. Each experiment with 8 aggregates.). Differences between samples were assessed by the Scheffe multiple comparison test; **, p < 0.01. The data are expressed as mean ±standard error. (B): The average number of CX26+ vesicles per aggregate from SB-treated (SB 1-10μM), and BMP4/SB (SB 1μM+BMP4 10ng/ml) treated aggregates. (n = 9 aggregates. For assessments, three time procedures were repeated to generate cells.). Statistical differences between samples were assessed by the One-way ANOVA and multiple comparison test: *, p < 0.05; **, p < 0.01. The data are expressed as mean ± the standard error.

**S1 Video. The three-dimensional (3D) image of whole CX26(+) vesicle in Day 7 aggregate.** The 3D image was reconstructed from consecutive slices of CX26(+) vesicle in (Figure 4A). Confocal stacks of CX26 (red), F-actin (green), and DAPI (blue) stain showing CX26-GJP-forming cells within the clear small vesicle.

**S2 Video. The three-dimensional (3D) image of CX26-GJP forming cells in CX26(+) vesicle.** The 3D image was reconstructed from consecutive slices of (Figure 4B). Confocal stacks of CX26 (red), F-actin (green), and DAPI (blue) stain showing that CX26 formed gap junction plaques at the cell-cell border.

